# Physical activity modifies the metabolic profile of CD4+ and CD8+ T cell subtypes at rest and upon activation in older adults

**DOI:** 10.1101/2024.08.20.607078

**Authors:** Edward Withnall, Jon Hazeldine, Alba Llibre, Niharika A. Duggal, Janet M. Lord, Amanda V. Sardeli

## Abstract

Aging reduces the functional competence of T cells. T cell metabolism regulates their function, with mitochondrial defects in mice resulting in aged phenotypes, including accelerated senescence. Physical activity (PA) maintains T cell function in older adults, although the mechanisms underlying this effect are poorly understood. This study examined the effects of aging on the metabolic profile of T cell subsets and investigated whether PA could improve metabolic function in T cells from older donors. We recruited nine young adults (23 ± 3y) and 19 healthy older adults who had high PA (HPA, N=9, 75.5 ± 4.7y) or low PA levels (LPA, N=10, 76.4 ± 2.1y), based on their moderate-to-vigorous PA scores. We investigated the metabolic profiles of CD4+ and CD8+ T cells at rest and post-activation (PMA and ionomycin), via SCENITH flow cytometry. Compared to young adults, older adults had higher mitochondrial dependence (MD) in unstimulated CD4+ and CD8+ naive, effector memory (EM) and central memory (CM); and higher protein synthesis in CD4+ EM, CD4+ naïve, CD8+ EM, suggesting higher energetic demand in T cells with aging. Upon activation there was a lower reduction in MD of CD4+ EMRA and CD8+ EMRA; and a greater increase in IL-6 and TNFα expression in CD8+ cells of older than young adults, indicative of impaired metabolic flexibility with aging. PA effects were more prominent in unstimulated CD8+ cells, where HPA had lower glucose dependence (GD) for overall CD8+, CD8+EM and a trend to higher MD in CD8+ CM than LPA. Upon activation, HPA had a lower increase in CD4+ TNFα expression and trended to have a higher reduction in MD of overall CD4+ and a higher reduction in GD of CD4+ EMRA, than LPA. This suggests a lower metabolic demand in CD4+ T cells of HPA. We concluded that PA could modify T cell metabolic profile at rest, and following activation, in older adults, which may explain the better T cell function in physically active older individuals.

---

Aging is associated with a progressive functional decline in the immune system (immunosenescence). Hallmarks of T cell ageing include: reduced naïve T cell counts, accumulation of terminally differentiated memory cells with senescent properties and a state of functional decline (proliferation, cytokine production and cytotoxicity) ultimately leads to an increased susceptibility to diseases and impaired response to vaccination and infections in older adults^1–3^. It is well known that physical activity (PA) exerts a beneficial impact on aged T cells, reflected in clinical outcomes such as better vaccination responses^4^ and reduced severity and duration of respiratory infections^5,6^, reducing age-related multimorbidity^7,8^. Previously, we have shown that maintenance of PA into older age can partially prevent immunosenescence, with PA preserving thymic output and reducing systemic inflammation^9^. However, the mechanisms by which PA preserves a youthful immune phenotype is poorly understood.

An important prerequisite for T cell function is their engagement in specific metabolic pathways that will define cell fate and function ^10^, and these pathways are known to be altered by aging^2,11^. PA is a potent metabolic regulator^12,13^. For example, Effector T cells undergo a rapid upregulation of their glycolytic metabolism (glycolytic shift)^2,11^, to generate biosynthetic intermediates, which are essential for increasing cell numbers after activation. Inhibition of glycolysis in activated T cells impedes effector cell differentiation, biasing cells towards memory subtypes that display increased oxidative metabolism and augmented mitochondrial fusion to sustain cellular longevity^2,11,14^. We therefore hypothesized that PA maintains T cell function by delaying metabolic dysfunction in older adults.

We recruited 9 young (23 ± 3 years; 7 males, 2 females) and 19 healthy older adults, subdivided based on their PA levels, as either high PA (HPA, moderate to vigorous [MVPA] 147 ± 18min/day; 76 ± 4 years; 4 males, 5 females) or low PA (LPA, MVPA 74 ± 27min/day; 72 ± 2 years; 4 males, 6 females). Peripheral blood mononuclear cells (PBMCs) were isolated, and the metabolic profiles of helper CD4^+^ and cytotoxic CD8^+^ T cells subtypes at rest, and post-activation (PMA [50 ng/ml] and Ionomycin [500 ng/ml]), examined by flow cytometry^15^.

Using SCENITH^15^, a flow cytometry based method for metabolic profiling, we assessed glucose dependence (GD) and mitochondrial dependence (MD), as a percentage of total protein synthesis. Recognizing the heterogeneity within T cells, we studied the metabolic profiles of naive (CCR7^+^ CD45RA^+^), central memory (CM, CCR7^-^ CD45RA^+^), effector memory (EM, CCR7^-^ CD45RA^-^) and terminally differentiated effector memory (EMRA, CCR7^-^ CD45RA^+^) CD4^+^ and CD8^+^ T cells, both at rest (unstimulated) and following activation. Detailed methods are provided (S1).

Our data showed that aging increased MD in all subtypes of CD4+ and CD8+ T cells at rest, excepting CD4^+^ EMRA and CD8^+^ EMRA (Figure 1). In parallel, advancing age reduced GD in CD8^+^ and there was a trend (p=0.067) to lower GD in CD4^+^ cells (S7). There was no difference in GD for any specific T cell subtype between young and old. Aged T cells with dysfunctional mitochondria display impaired respiration, reduced glycolysis and inefficient one-carbon metabolism^1,2,16^. Our data suggest that older adults may sustain higher mitochondria dependence to compensate for impaired glucose metabolism. In fact, previous studies showed that older individuals have increased mitochondrial mass and fusion, but also defective autophagy and increased influx and storage of lipids, which impairs T cell proliferation and immune responses in older adults^11,17–20^. Interestingly, we saw higher protein synthesis (total puromycin) for CD4+ EM, CD4+ naïve, CD8+ EM in older adults, compared to young, suggesting they need higher energy demand to sustain baseline functions (S5). This could be due to the chronic activation of the immune system that occurs with aging^21^, as well as cellular senescence, mitochondrial dysfunction, reduced autophagy and higher prevalence of chronic infections such as cytomegalovirus^22^.

**Figure 1.**
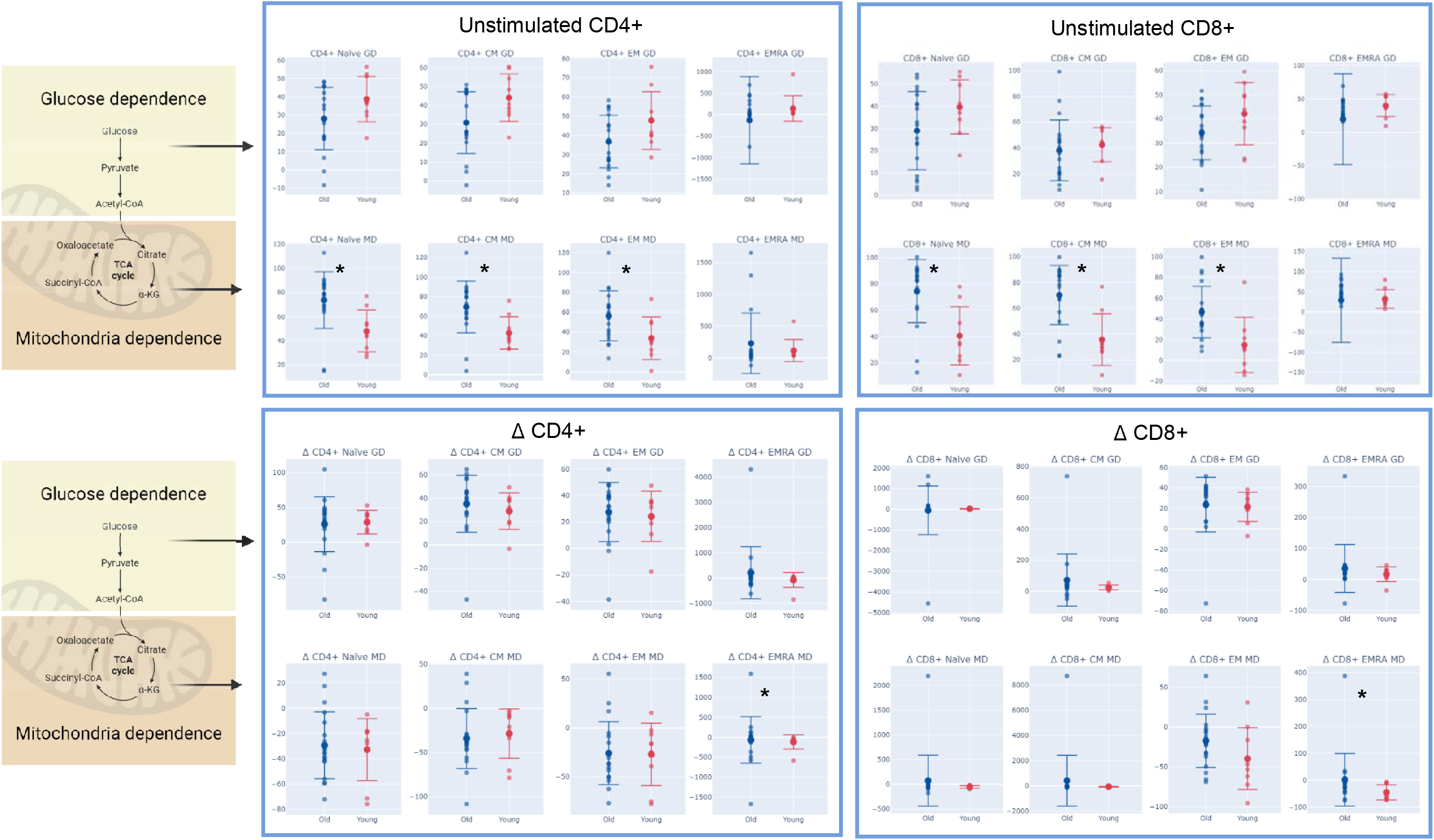
Effects of aging on the metabolic profile of T cells. Legend: GD: Glucose dependence (%); MD: mitochondria dependence (%); Δ: change with stimulation (stimulated minus unstimulated); *: significant difference between age groups (p-value ≤ 0.05).

Upon activation, young subjects had a higher reduction in MD in CD8^+^ T cells and a trend (p=0.079) to higher increase in GD in CD4+ (S7), compared to older adults, reinforcing their more efficient metabolic shift. Specifically, these effects were led by EMRA, which showed a significantly higher reduction in MD of CD4^+^ EMRA and CD8^+^ EMRA (Figure 1B), and that this T cell population was more prevalent in older adults (S4). It is well known that naïve, aged and senescent T cells of old mice and humans have an impaired switch to glycolytic metabolism upon activation (reduced metabolic flexibility), compared to young individuals^2,11,17,23,24^. Those cells show increased fatty acid uptake and lipid storage after stimulation^17^. The accumulation of glucose and lipids in healthy CD8+ cells, contributes to a senescent profile, with increased p-p53 expression and a fragmented mitochondrial morphology, similar to CD8^+^ EMRA of diabetic older adults^25^. Therefore, it is possible the sustained higher mitochondria dependence in older adults, is associated to the accumulation of these intracellular substrates favoring T cell exhaustion^26^.

Intracellular cytokine expression was very similar between groups. Nonetheless, even with lower metabolic flexibility, older adults had a higher percentage of CD8^+^ T cells producing IL-6 (Δ young: 12.4 ± 26.7 vs. Δ Old: 21.1 ± 22.3, p=0.042) and a trend (p=0.067) to have higher TNFα in CD8^+^ than young upon activation (S6). A robust increase in IL-6 and TNFα is necessary for an efficient immune response in T cells^27–29^. However, in agreement with our data, a higher inflammatory phenotype has been observed after short term activation in older adults^30–33^. This higher pro-inflammatory cytokine production could be due to the higher frequency of senescent cells in older adults ^34^. Although we did not specifically assess senescent cells here, older adults had higher frequency of CD8^+^ EMRAs, which encompass the senescent-like T cells^35^ (S4). In any case, the higher protein synthesis, in CD4+ and CD8+ subtypes upon stimulation in older adults (S5), suggests a higher demand in T cells of older adults compared to young adults.

In older adults, different PA levels did not lead to significant differences in unstimulated CD4^+^ metabolic profile (Figure 2). HPA group had lower GD in unstimulated CD8^+^, CD8^+^EM and CD4^+^EM, than LPA. We saw earlier that aging increased MD in most CD4^+^ and CD8^+^ subtypes, except EMRA, and reduced GD in CD8^+^ cells (S7). Therefore, this data suggests PA did not overcome the aging effects in the metabolism of T cells at rest. Instead, HPA had a trend (p=0.095) to an additional increase in MD in CD8^+^ CM, which might be related to physical fitness improving mitochondrial respiration in T cells^36,37^. Others have reported that an increase in AMPK activity with exercise, favors beta oxidation in T cells and reduces lipid accumulation, ameliorating immunosenescence^25,38,39^.

**Figure 2.**
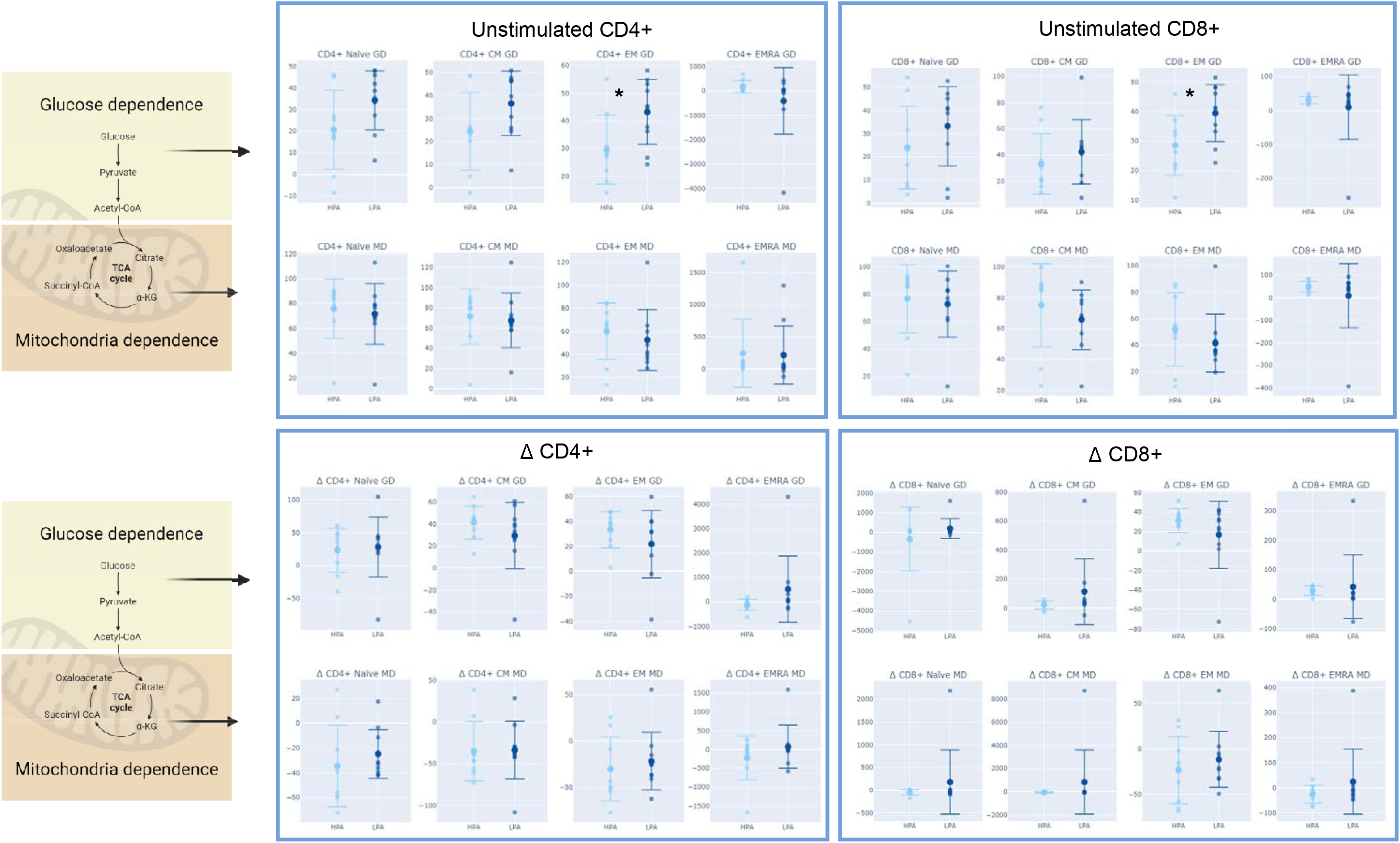
Effects of physical activity on the metabolic profile of T cells. Legend: GD: Glucose dependence (%); MD: mitochondria dependence (%); Δ: ch with stimulation (stimulated minus unstimulated); *: significant difference between age groups (p-value ≤ 0.05).

Following activation (Figure 2B), HPA adults trended (p=0.079) towards higher reduction in MD in overall CD4^+^ (S7) and a lower increase in GD of CD4^+^ EMRA (p=0.065), than LPA adults. We also observed significantly lower TNFα expression in CD4^+^ cells of HPA, compared to LPA (S6). Although an increase in glucose uptake is required for T cell activation, excessive glucose uptake can be detrimental^40,41^; and inhibiting glycolytic metabolism can enhance the function of certain T cell subtypes such as CD8^+^ T cell memory^14^. Furthermore, this lower metabolic shift can be simply due to lower energetic demand in physically active individuals. In fact, in other cell types, such as skeletal muscle, exercise is known to improve insulin sensitivity, glucose uptake, lactate clearance, increasing systemic metabolic flexibility^13^.

It is noteworthy that this trend towards reduced metabolic demand in HPA was accompanied by lower TNF production in CD4^+^ T cells (S5), thus demonstrating that PA can counteract the higher inflammatory phenotype that has been observed after short term activation in older adults^30–33^. It is not clear, why in our study, we did not observe an increase in cytokine production by CD4^+^ cells with age, as we had observed for CD8^+^, but this could be due to the lack of power in our analysis. In any case, a reduced need for high levels of cytokine production in HPA might be associated with improved cytokine sensitivity that is known to occur with PA^42–44^.

In summary, advancing age increased MD in CD4^+^ and CD8^+^ naïve, EM and CM at rest, and impaired metabolic flexibility in CD4^+^ and CD8^+^ EMRA upon activation. Interestingly, in older adults, PA led to even higher MD in baseline CD8^+^ cells and trended to improve metabolic flexibility upon activation in CD4^+^ cells. There was higher cytokine production and metabolic demand in CD8^+^ cells with aging; however, PA seemed to reduce the energetic demand and cytokine production only in CD4^+^ cells upon activation. A limitation of our study was the small sample size, given the high heterogeneity that is seen in older adults. Therefore, we believe that future analysis with larger sample sizes may reveal more significant effects of physical activity on T cell metabolism. Also, we used PMA and Ionomycin to activate T cells, and a more physiological stimulus, such as anti-CD3 and anti-CD28, could lead to a clearer separation within the different T cell subtypes.

In conclusion, we identified for the first time that PA modified the metabolic profile in a variety of T cell subtypes, in older adults, at rest and following activation. The effects can partially explain the better T cell function seen in physically active individuals. If this is confirmed, future research in this field should aim to identify the cellular regulators of T cell fitness with PA.

S1 to S8 can be accessed in the Supplementary information.

## Abbreviations

CM: Central memory
EM: Effector memory
EMRA: Effector memory re-expressing CD45RA
GD: Glucose dependence
HPA: High physical activity level group Interleukin 6
IL-6: Interleukin6
IL-17: Interleukin 17
LPA: Low physical activity level group
MD: Mitochondria dependence
MVPA: Moderate to vigorous physical activity
PA: Physical activity
TNF: Tumour necrosis factor alpha
IFN: Interferon gamma

## AUTHOR CONTRIBUTIONS

AVS conceived the study, secured funding, and managed overall direction. AVS and JH designed experiments. AVS and ED collected samples. ED performed all experiments. EW and AVS analyzed data and write first draft of the manuscript. JH, ND, AL and JL critically revised data interpretation and provided critical feedback that helped shape the final manuscript.

## ACKNOWLEDGMENTS

The authors would like to thank Dr. Johannes Schroth and professor Sian Henson for their expertise and assistance with the optimization of the SCENITH methods.

## References

1. Desdín-Micó G, Soto-Heredero G, Aranda JF, Oller J, Carrasco E, Gabandé-Rodríguez E, et al. T cells with dysfunctional mitochondria induce multimorbidity and premature senescence. Science. 2020; 368(6497):1371–6.

2. Møller SH, Hsueh PC, Yu YR, Zhang L, Ho PC. Metabolic programs tailor T cell immunity in viral infection, cancer, and aging. Cell Metab. 2022;34(3):378–95.

3. Pinti M, Appay V, Campisi J, Frasca D, Fülöp T, Sauce D, et al. Aging of the immune system: Focus on inflammation and vaccination. Eur J Immunol. 2016;46(10):2286– 301.

4. Pascoe AR, Fiatarone Singh MA, Edwards KM. The effects of exercise on vaccination responses: A review of chronic and acute exercise interventions in humans . Vol. 39, Brain, Behavior, and Immunity. Academic Press Inc.; 2014; p. 33–41.

5. Grande AJ, Keogh J, Silva V, Scott AM. Exercise versus no exercise for the occurrence, severity, and duration of acute respiratory infections . Vol. 2020, Cochrane Database of Systematic Reviews. 2020.

6. DC N, DA H, MD A, Sha W. Upper respiratory tract infection is reduced in physically fit and active adults. Br J Sports Med . 2011;45(12):987–92.

7. Sallis R, Young DR, Tartof SY, Sallis JF, Sall J, Li Q, et al. Physical inactivity is associated with a higher risk for severe COVID-19 outcomes: a study in 48 440 adult patients. Br J Sports Med. 2021;55(19):1099–105.

8. Duggal NA, Niemiro G, Harridge SDR, Simpson RJ, Lord JM. Can physical activity ameliorate immunosenescence and thereby reduce age-related multi-morbidity? . Vol. 19, Nature Reviews Immunology. 2019; p. 563–72.

9. Duggal NA, Pollock RD, Lazarus NR, Harridge S, Lord JM. Major features of immunesenescence, including reduced thymic output, are ameliorated by high levels of physical activity in adulthood. Aging Cell. 2018; 17(2).

10. Voss K, Hong HS, Bader JE, Sugiura A, Lyssiotis CA, Rathmell JC. A guide to interrogating immunometabolism. Nat Rev Immunol. 2021;21(10):637–52.

11. Quinn KM, Vicencio DM, La Gruta NL. The paradox of aging: Aging-related shifts in T cell function and metabolism. Semin Immunol. 2023;70.

12. Hodgman CF, Hunt RM, Crane JC, Elzayat MT, LaVoy EC. A Scoping Review on the Effects of Physical Exercise and Fitness on Peripheral Leukocyte Energy Metabolism in Humans. Exerc Immunol Rev. 2023;29:54–87.

13. Smith JAB, Murach KA, Dyar KA, Zierath JR. Exercise metabolism and adaptation in skeletal muscle. Nat Rev Mol Cell Biol. 2023;24(9):607–32.

14. Sukumar M, Liu J, Ji Y, Subramanian M, Crompton JG, Yu Z, et al. Inhibiting glycolytic metabolism enhances CD8+ T cell memory and antitumor function. J Clin Invest. 2013;123(10):4479.

15. Argüello RJ, Combes AJ, Char R, Gigan JP, Baaziz AI, Bousiquot E, et al. SCENITH: A Flow Cytometry-Based Method to Functionally Profile Energy Metabolism with Single-Cell Resolution. Cell Metab. 2020;32(6):1063-1075.e7.

16. Ron-Harel N, Notarangelo G, Ghergurovich JM, Paulo JA, Sage PT, Santos D, et al. Defective respiration and one-carbon metabolism contribute to impaired naïve T cell activation in aged mice. Proc Natl Acad Sci U S A. 2018;115(52):13347–52.

17. Nicoli F, Cabral-Piccin MP, Papagno L, Gallerani E, Fusaro M, Folcher V, et al. Altered Basal Lipid Metabolism Underlies the Functional Impairment of Naive CD8+ T Cells in Elderly Humans. J Immunol. 2022;208(3):562–70.

18. Bektas A, Schurman SH, Gonzalez-Freire M, Dunn CA, Singh AK, Macian F, et al. Age-associated changes in human CD4+ T cells point to mitochondrial dysfunction consequent to impaired autophagy. Aging (Albany NY). 2019;11(21):9234–63.

19. Alsaleh G, Panse I, Swadling L, Zhang H, Richter FC, Meyer A, et al. Autophagy in T cells from aged donors is maintained by spermidine and correlates with function and vaccine responses. Elife. 2020;9:1–21.

20. Jin J, Li X, Hu B, Kim C, Cao W, Zhang H, et al. FOXO1 deficiency impairs proteostasis in aged T cells. Sci Adv. 2020; 6(17).

21. Franceschi C, Bonafè M, Valensin S, Olivieri F, De Luca M, Ottaviani E, et al. Inflamm-aging. An evolutionary perspective on immunosenescence. Ann N Y Acad Sci. 2000;908:244–54.

22. Haynes L. Aging of the Immune System: Research Challenges to Enhance the Health Span of Older Adults. Front Aging. 2020;1.

23. Ron-Harel N, Notarangelo G, Ghergurovich JM, Paulo JA, Sage PT, Santos D, et al. Defective respiration and one-carbon metabolism contribute to impaired naïve T cell activation in aged mice. Proc Natl Acad Sci U S A. 2018;115(52):13347–52.

24. Nian Y, Iske J, Maenosono R, Minami K, Heinbokel T, Quante M, et al. Targeting age-specific changes in CD4+ T cell metabolism ameliorates alloimmune responses and prolongs graft survival. Aging Cell. 2021; 20(2).

25. Callender LA, Carroll EC, Garrod-Ketchley C, Schroth J, Bystrom J, Berryman V, et al. Altered Nutrient Uptake Causes Mitochondrial Dysfunction in Senescent CD8+ EMRA T Cells During Type 2 Diabetes. Front aging. 2021;2.

26. Yu YR, Imrichova H, Wang H, Chao T, Xiao Z, Gao M, et al. Disturbed mitochondrial dynamics in CD8+ TILs reinforce T cell exhaustion. Nat Immunol. 2020;21(12):1540– 51.

27. Tanaka T, Narazaki M, Kishimoto T. IL-6 in inflammation, immunity, and disease. Cold Spring Harb Perspect Biol. 2014; 6(10).

28. Sedger LM, McDermott MF. TNF and TNF-receptors: From mediators of cell death and inflammation to therapeutic giants - past, present and future. Cytokine Growth Factor Rev. 2014;25(4):453–72.

29. Dong C. Cytokine Regulation and Function in T Cells. Annu Rev Immunol. 2021;39:51–76.

30. Akbar AN, Henson SM, Lanna A. Senescence of T Lymphocytes: Implications for Enhancing Human Immunity. Trends Immunol. 2016;37(12):866–76.

31. McNerlan SE, Rea IM, Alexander HD. A whole blood method for measurement of intracellular TNF-α, IFN-γ and IL-2 expression in stimulated CD3 + lymphocytes: Differences between young and elderly subjects. Exp Gerontol. 2002; 37(2–3):227–34.

32. O’Mahony L, Holland J, Jackson J, Feighery C, Hennessy TPJ, Mealy K. Quantitative intracellular cytokine measurement: age-related changes in proinflammatory cytokine production. Clin Exp Immunol. 1998;113(2):213–9.

33. Henson SM, Macaulay R, Riddell NE, Nunn CJ, Akbar AN. Blockade of PD-1 or p38 MAP kinase signaling enhances senescent human CD8+ T-cell proliferation by distinct pathways. Eur J Immunol. 2015;45(5):1441–51.

34. Freund A, Orjalo A V., Desprez PY, Campisi J. Inflammatory Networks during Cellular Senescence: Causes and Consequences. Trends Mol Med. 2010;16(5):238.

35. Callender LA, Carroll EC, Beal RWJ, Chambers ES, Nourshargh S, Akbar AN, et al. Human CD8 + EMRA T cells display a senescence-associated secretory phenotype regulated by p38 MAPK. Aging Cell. 2018; 17(1).

36. Gebhardt K, Hebecker A, Honekamp C, Nolte S, Barthkuhn M, Wilhelm J, et al. Respiratory and Metabolic Responses of CD4 + T Cells to Acute Exercise and their Association with Cardiorespiratory Fitness. Med Sci Sports Exerc. 2024.

37. Andonian B, Bartlett D, Koss A, Muoio D, Koves T, Clair ES, et al. High-Intensity Interval Training Increases Rheumatoid Arthritis Cardiorespiratory Fitness in Association with Improvements in CD4+ T Cell and Skeletal Muscle Mitochondrial Metabolism. ACR Converg. 2020.

38. Rosa-Neto JC, Lira FS, Little JP, Landells G, Islam H, Chazaud B, et al. Immunometabolism-fit: How exercise and training can modify T cell and macrophage metabolism in health and disease. Exerc Immunol Rev. 2022;28:29–46.

39. Padilha CS, Figueiredo C, Minuzzi LG, Chimin P, Deminice R, Krüger K, et al. Immunometabolic responses according to physical fitness status and lifelong exercise during aging: New roads for exercise immunology. Ageing Res Rev. 2021.

40. Chen DL, Wang X, Yamamoto S, Carpenter D, Engle JT, Li W, et al. Increased T cell glucose uptake reflects acute rejection in lung grafts. Am J Transplant. 2013; 13(10):2540–9.

41. Sengupta S, Vitale RJ, Chilton PM, Mitchell TC. Adjuvant induced glucose uptake by activated T cells is not correlated with increased survival. Adv Exp Med Biol. 2008;614:65–72.

42. Nash D, Hughes MG, Butcher L, Aicheler R, Smith P, Cullen T, et al. ILLJ6 signaling in acute exercise and chronic training: Potential consequences for health and athletic performance. Scand J Med Sci Sports. 2023 Jan 1 [cited 2024 Aug 7];33(1):4.

43. Keller C, Steensberg A, Hansen AK, Fischer CP, Plomgaard P, Pedersen BK. Effect of exercise, training, and glycogen availability on IL-6 receptor expression in human skeletal muscle. J Appl Physiol. 2005 Dec;99(6):2075–9.

44. Hunter CA, Jones SA. IL-6 as a keystone cytokine in health and disease. Nat Immunol. 2015;16(5):448–57.

