## Supplementary information for "Physical activity modifies the metabolic profile of CD4+ and CD8+ T cell subtypes at rest and upon activation in older adults"

**Supporting Information**

**S1. Methods.**

***Population.*** Ethical approval for this study was granted by the North West - Haydock Research Ethics Committee (Reference: 22/NW/1087) and written consent was obtained from individuals before participation.

Older participants had to be above 65 years of age and younger participants had to be between 18-35 years of age. All participants were required to be in good health at the time of the study (based on the criteria set out by Greig *et al* ^1^), this was defined as without infection, disease, or taking immunosuppressive medication. Results from individuals with cancer, chronic inflammatory conditions, or taking immunosuppressive medication were excluded prior to data analysis. Participants completed a health questionnaire, a physical functioning assessment, and donated blood samples. The physical functioning assessment consisted of hand grip strength, walking gait speed, and 30 second sit to stand tests. Blood samples were collected into heparinised vacutainers® (BD Biosciences, New Jersey, USA).

***PBMC isolation.*** The collected blood samples were pooled from heparinised vacutainers into clean 25ml universal tubes (Scientific Laboratory Supplies, Nottingham, UK) and diluted with GPS only RPMI media (Life Technologies Limited, Paisley, UK) at a 1:1 ratio. The blood-RPMI mix was layered on top of 6mls of Ficoll-Plaque^TM^ Plus (GE Healthcare, Buckinghamshire, UK) and centrifuged at 400 × g for 30 minutes at room temperature (RT) with no break or acceleration. After centrifugation, the peripheral blood mononuclear cell (PBMC) layer was transferred via Pasteur pipette (Scientific Laboratory Supplies, Nottingham, UK) into a fresh universal tube containing autoMACS running buffer (Miltenyi Biotec, Surrey, UK). Once all the PBMCs were transferred, the universal tube was topped up with autoMACS running buffer and centrifuged at 300 × g for 10 mins at RT with full break and acceleration. Post spin, the pellet was resuspended in 25ml autoMACS running buffer and washed via centrifugation at 300 × g for 10 minutes at RT. 1ml of prepared freezing solution containing 5ml 10% DMSO (Sigma Alrich, Dorset, UK) and 45 ml heat inactivated FCS (Thermofisher, Massachusetts, USA) was used to resuspend the pellet. 500µl of PBMCs were aliquoted into 1ml Cryo.s (Greiner Bio-One, Gloucestershire, UK) and then placed in a RT Mr Frosty^TM^ Freezing Container (ThermoFisher, Massachusetts, USA) in which they were frozen at -80°C.

***PBMC stimulation.*** PBMCs were removed from the freezer, thawed at 37°C and added to a universal tube containing 10ml of warm RPMI medium with no penicillin andstreptomycin. This PBMC-RMPI medium mix was centrifuged at 300 × g for 10 minutes at room temperature (RT), resuspended in 1ml of warm RMPI medium with no P/S, and incubated for 15 hours at 37°C, 5% CO_2_. Post incubation, 9ml of warm RPMI medium with no P/S was added to the PBMCs and the mix was centrifuged at 300 × g for 10 minutes at RT and resuspended in 1ml of warm RPMI medium with no penicillin and streptomycin. 100µl of this mix were added to a 1.5ml microtube (Sarstedt AG & Co., Nümbrecht, Germany) and the white blood cell (WBC) count was determined via Sysmex (Sysmex, Milton Keynes, UK). The PBMC mixture was diluted with warm RPMI medium with no penicillin and streptomycin to obtain a concentration of 2×10^6^ WBC/ml. 100µl of the PBMC mixture were aliquoted to 5ml polypropylene round-bottom tubes (Scientific Laboratory Supplies, Nottingham, UK), stimulated with 2µl Phorbel 12-myristate 13-acetate (PMA) (50ng/ml; Merck Life Science, Dorset, UK) and 2µl Ionomycin (500ng/ml; Merck Life Science, Dorset, UK), and incubated at 37°C, 5% CO_2_. Untreated PBMCs were used as negative controls.

***T cell metabolic profile.*** A new flow cytometry method, single cell energetic metabolism by profiling translation inhibition or SCENITH ^2^, was used to investigate the metabolic profiles of CD4^+^ and CD8^+^ T cell subtypes. Almost half of the energy produced by mammalian cells via metabolic processes is used up by protein synthesis Machinery^3^. The incorporation of puromycin is a widely accepted readout for protein synthesis in vitro and in vivo^4–6^.Therefore, using an anti-puro monoclonal antibody, the metabolic activity of cells can be measured by puromycin levels via flow cytometry. Metabolic inhibitors are utilized to determine the metabolic profile of a cell, defined by glucose and mitochondria dependence. The 2-Deoxy-D-Glucose (2DG) inhibit glycolysis and therefore allows the calculation of glucose dependence^7^, while Oligomycin inhibits oxidative phosphorylation, allowing calculation of mitochondria dependence^8^.

Immediately after being stimulated or left untreated as the control, PBMCs were treated with either 10µl of 2DG (100mM; Merck Life Science, Dorset, UK), 1µl Oligomycin from *Streptomyces diastatochromogenes* (1µM; Merck Life Science, Dorset, UK]), a sequential combination of the drugs at the same concentrations, or left untreated as the control, and incubated at 37°C, 5% CO_2_ for 15 minutes. Following incubation, PBMCs were treated with 5µl Puromycin dihydrochloride from *Streptomyces alboniger* (10µg/ml; Merck Life Science, Dorset, UK) and left to incubate at 37°C, 5% CO_2_ for a further 30 minutes. PBMCs were then washed with 200µl phosphate buffered saline (PBS) via centrifugation at 250 × g for 5 minutes at 4°C and stained with surface marker conjugated antibodies CD3 (PE-Cyanine7; Life Technologies Corp, California, USA), CD4 (Brilliant Violet 421^TM^; BioLegend®, California, USA), CD8 (PE; BioLegend®, California, USA), CCR7 (APC; BioLegend®, California, USA), and CD45RA (PerCP; BioLegend®, California, USA) and left to incubate on ice for 20 minutes protected from light. Post incubation, PBMCs were washed with 300µl PBS at 250 × g for 5 minutes at 4°C and then fixed and permeabilised (FOXP3 transcription factor Staining Buffer Set; Fisher Scientific, Leicestershire, UK), stained with Human TruStain FcX^TM^ (BioLegend®, California, USA) and anti-Puromycin (Alexa Fluor® 488 anti-puromycin; BioLegend®, California, USA) or isotype control (Alexa Fluor® 488, and incubated for an hour at 4°C without any light. After staining, PBMCs were washed with 200µl PBS at 250 × g for 5 minutes at 4°C and resuspended in 200µl PBS. Samples were kept on ice and protected from light prior to analysis.

***T cell cytokine production.*** After a 4-hour incubation period with Brefeldin A (10µg/ml; Merck Life Science, Dorset, UK) and with or without stimulation, PBMCs were washed with 300µl PBS via centrifugation at 250 × g for 5 minutes at 4°C. Once resuspended in 50µl PBS, PBMCs were stained with surface marker conjugated antibodies CD3 (PE-Cyanine7; Life Technologies Corp, California, USA), CD4 (Brilliant Violet 421^TM^; BioLegend®, California, USA), CD8 (VioGreen^TM^; Miltenyi Biotec, Surrey, UK), CCR7 (APC; BioLegend®, California, USA), and CD45RA (PerCP; BioLegend®, California, USA) and left to incubate in ice protected from light for 20 minutes. Post incubation PBMCs were washed with 300µl PBS at 250 × g for 5 minutes at 4°C and fixed via a 30-minute incubation with 50µl Medium A (Life Technologies LTD, Paisley, UK) at RT. PBMCs were then washed via centrifugation at 250 × g for 5 minutes at 4°C and resuspended in 50µl Medium B for permeabilization. PBMCs underwent intracellular staining for IL-6 (PE; BioLegend®, California, USA) and TNFα (FTIC; BD Pharmingen, California, USA) and were then left to incubate on ice for 30 minutes without light. After incubation, PBMCs were washed with 300µl PBS at 250 × g for 5 minutes at 4°C and resuspended in 200µl PBS. Samples were kept on ice and protected from light prior to analysis.

***Flow cytometry.*** For the SCENITH analysis the distribution of CD4^+^ and CD8^+^ T cells subsets were determined. Each T cell subset were defined by CD4^+^ naïve (CD3^+^CD4^+^CD45RA^+^CCR7^+^), CD4^+^ central memory (CM, CD3^+^CD4^+^CD45RA^-^CCR7^+^), CD4^+^ effector memory (EM, CD3^+^CD4^+^CD45RA^-^CCR7^-^), CD4^+^ terminally differentiated effector memory cells (CD3^+^CD4^+^CD45RA^+^CCR7^-^), CD8^+^ naïve (CD3^+^CD8^+^CD45RA^+^CCR7^+^), CD8^+^ central memory (CM, CD3^+^CD8^+^CD45RA^-^CCR7^+^), CD8^+^ effector memory (EM, CD3^+^CD8^+^CD45RA^-^CCR7^-^), CD8^+^ terminally differentiated effector memory cells (CD3^+^CD8^+^CD45RA^+^CCR7^-^) (S2 a-d). The metabolic profile of each T cell subset was quantified by the median fluorescence intensity (MFI) of puromycin post treatment with metabolic inhibitors (2DG and Oligomycin), isotype control (S2 e and f). Metabolic profile was defined by glucose and mitochondria dependence, and we also considered maximum protein synthesis as Puromycin MFI (without inhibition) Glucose and mitochondria dependence were determined by the calculations provided:

$$Glucose dependence=\frac{100(PuroMFI-PuroMFI with 2DG inhibition)}{(PuroMFI-PuroMFI with 2DG and Oligomycin inhibition)}$$

$$Mitochondria dependence=\frac{100(PuroMFI-PuroMFI with Oligomycin inhibiton)}{(PuroMFI-PuroMFI with 2DG and Oligomycin inhibition}$$

For the T cell cytokine experiments, CD4^+^ and CD8^+^ T cell populations were determined. These populations were defined by: CD4^+^ (CD3^+^CD4^+^CD8^-^), CD8^+^ (CD3^+^CD4^-^CD8^+^) (S3 a-b). To access the cytokine production of the CD4^+^ and CD8^+^ T cell populations, TNFα and IL-6 MFI and % were measured via anti-TNF and anti-IL-6 (S3 c-f).

All flow cytometry was performed on a MACSQuant® Analyzer 8 flow cytometer (Miltenyi Biotec. Surrey, UK) and analysed by FlowJo software (FlowJo LLC, Orlando, USA). The key resources are listed below (S8).

***Statistical analysis.*** Statistical analyses were performed using IBM SPSS software version 29.0 (IBM, Portsmouth, UK). The normality of the data was assessed using the Kolmogorov-Smirnov test. Majority of variables were non-normally distributed data, and therefore we used an Independent-Samples Mann-Whitney U test to compare the distributions between groups in each condition. Differences were considered statistically significant at p-value ≤ 0.05. Results were presented using GraphPad PRISM® software (GraphPad software, California, USA). Individual data with median and interquartile range were presented in the graphs, while baseline characteristics’ data were presented as mean ± standard deviation in the table and in the text.

**References**

1.	Greig CA, Young A, Skelton DA, Pippet E, Butler FMM, Mahmud SM. Exercise studies with elderly volunteers. Age Ageing. 1994 Nov 18];23(3):185–9.

2.	Argüello RJ, Combes AJ, Char R, Gigan J-P, Baaziz AI, Bousiquot E, et al. SCENITH: A Flow Cytometry-Based Method to Functionally Profile Energy Metabolism with Single-Cell Resolution. Cell Metab. 2020;32(6):1063-1075.e7.

3.	Lindqvist LM, Tandoc K, Topisirovic I, Furic L. Cross-talk between protein synthesis, energy metabolism and autophagy in cancer. Curr Opin Genet Dev. 2018;48:104–11.

4.	Aviner R. The science of puromycin: From studies of ribosome function to applications in biotechnology. Comput Struct Biotechnol J. 2020;18:1074–83.

5.	Hidalgo San Jose L, Signer RAJ. Cell-type-specific quantification of protein synthesis in vivo. Nat Protoc. 2019;14(2):441–60.

6.	Seedhom MO, Hickman HD, Wei J, David A, Yewdell JW. Protein Translation Activity: A New Measure of Host Immune Cell Activation. J Immunol. 2016;197(4):1498–506.

7.	Singh R, Gupta V, Kumar A, Singh K. 2-Deoxy-D-Glucose: A Novel Pharmacological Agent for Killing Hypoxic Tumor Cells, Oxygen Dependence-Lowering in Covid-19, and Other Pharmacological Activities. Adv Pharmacol Pharm Sci. 2023.

8.	Mackieh R, Al-Bakkar N, Kfoury M, Roufayel R, Sabatier JM, Fajloun Z. Inhibitors of ATP Synthase as New Antibacterial Candidates. Antibiot (Basel, Switzerland). 2023;12(4).

**S2. Gating strategy for SCENITH protocol.**

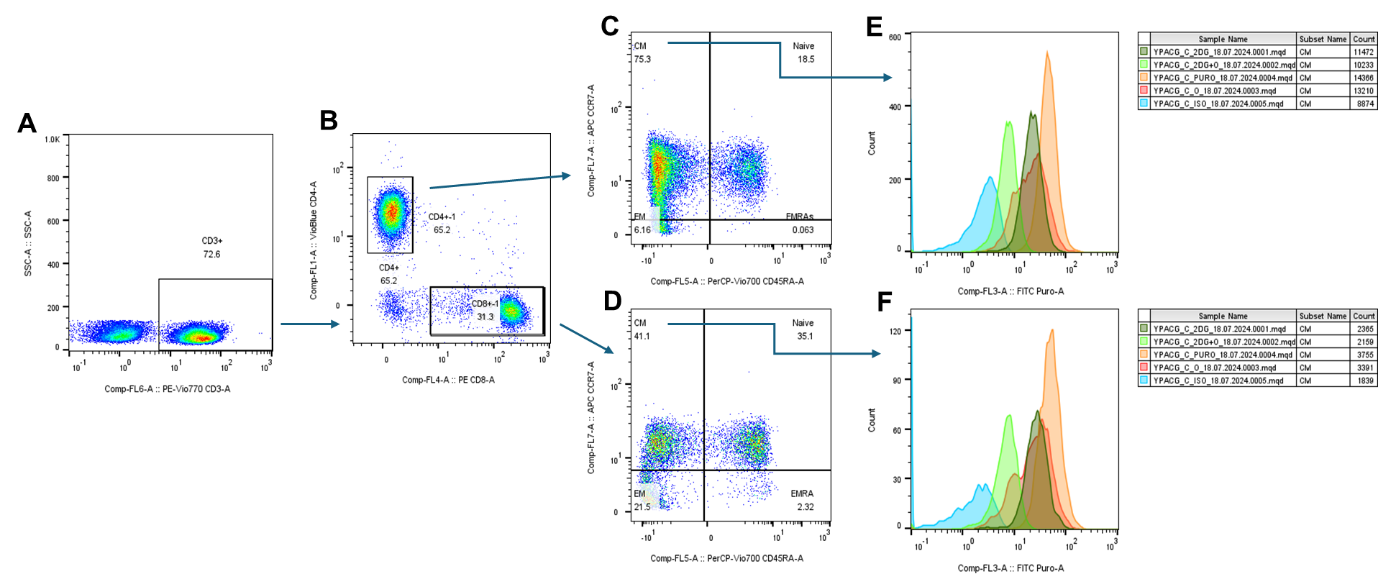

**Legend:** PBMCs were stained with anti-CD3 (A), anti-CD4 and anti-CD8 (B), anti-CD-45RA and anti-CCR7 (C and D), and Alexa Fluor 488 anti-puromycin and Alexa Fluor 488 Mouse IgG2a k isotype control (E and F).

**S3. Gating strategy for T cell cytokine production experiments.**

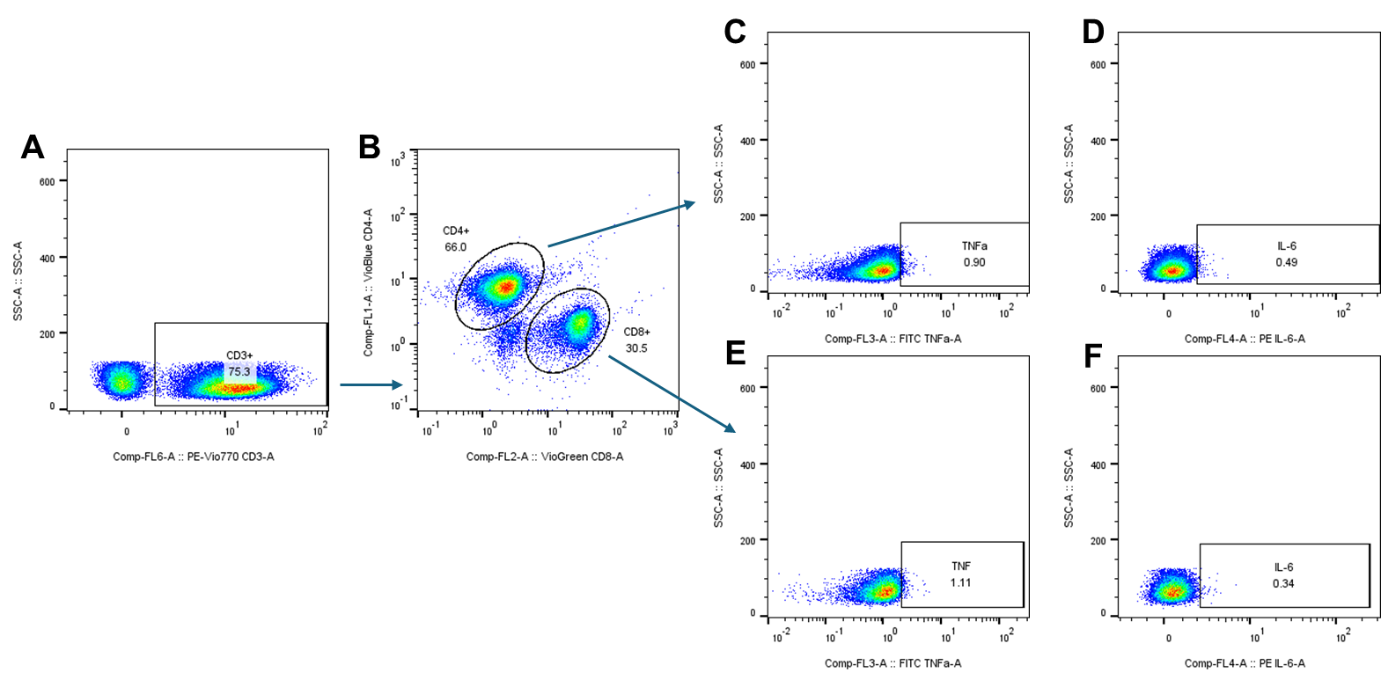

**Legend:** PBMCs were stained with anti-CD3 (A), anti-CD4 and anti-CD8 (B), and anti-TNFα and anti-IL-6 (C-F).

| **S4.** Participant characteristics and immune phenotype. | | | |
| --- | --- | --- | --- |
| ***General characteristics*** | **Young (n=9)** | **Old HPA (n=9)** | **Old LPA (n=10)** |
| Age (yrs) | **23.7 ± 3.2^bc^** | **75.6 ± 4.3^a^** | **76 ± 2^a^** |
| Sex | m=7, f=2 | m=4, f=5 | m=4, m=6 |
| Weight (kg) | 73.8 ± 16.9 | 66.4 ± 10.7 | 72.6 ± 20.1 |
| BMI (kg/m^2^) | 22.5 ± 3.4 | 24 ± 3 | 26.9 ± 5.1 |
| ***Physical activity and physical fitness*** |  |  |  |
| Mild (hours /week) | **5.7 ± 1.6^b^** | **9.6 ± 2.5^ac^** | **5.6 ± 3.8^b^** |
| Moderate (hours /week) | 2.6 ± 2.3 | 2.3 ± 1.5 | 0.7 ± 1.4 |
| Vigorous (hours/week) | **4.3 ± 2.8^c^** | 1.4 ± 1.7 | **0.9 ± 2.5^a^** |
| Sedentary time (hours/d) | 1.3 ± 1.3 | 0.7 ± 0.8 | 1.1 ± 1 |
| Grip Strength (kg) | **37.1 ± 9.8^c^** | 26.8 ± 6.6 | **23.6 ± 6.5^a^** |
| Gait Speed (m/s) | 1.3 ± 0.3 | 1.3 ± 0.3 | 1.2 ± 0.4 |
| 30-second sit-to-stand test (reps) | **19.8 ± 5^c^** | 13.9 ± 3.7 | **13.1 ± 4.4^a^** |
| **Immune phenotype** |  |  |  |
| T CD4+ % | 59.2 ± 9.2c | 69 ± 0.2 | 75.4 ± 13.8a |
| T CD4+ naïve % | 46.9 ± 16.5 | 40.8 ± 14.9 | 36.2 ± 19.1 |
| T CD4+ Central memory % | 40.1 ± 14.8 | 50.1 ± 11.7 | 51.1 ± 21.4 |
| T CD4+ Effector memory % | 12.1 ± 6.8 | 8.5 ± 5.1 | 9.7 ± 10.4 |
| T CD4+ EMRA % | 0.8 ± 0.9 | 0.6 ± 0.3 | 3 ± 6.3 |
| T CD8+ % | 34.5 ± 7.4c | 26.5 ± 15.3 | 20 ± 12.1a |
| T CD8+ naïve % | 45.1 ± 16.7c | 19.4 ± 11.3 | 17.3 ± 14.4a |
| T CD8+ Central memory % | 18.8 ± 15.8 | 6.5 ± 3.8 | 8.9 ± 7.7 |
| T CD8+ Effector memory % | 26.8 ± 13.8bc | 52.5 ± 12.9a | 51 ± 19.2a |
| T CD8+ EMRA % | 9.4 ± 7.9b | 21.7 ± 8.1a | 22.8 ± 20.5 |

**Legend:** Groups were defined by age and moderate-to-vigorous activity (MVPA). Participant characteristics were determined via a health questionnaire. The physical activity of participants was quantified via a health questionnaire and physical functioning assessment, consisting of hand grip strength, walking gait speed, and a 30-second sit-to-stand test. *The immune phenotype of participants was defined by PBMCs isolated from blood samples and assessed via flow cytometry. T cell subsets were defined as T CD4^+^: (CD3^+^CD4^+^), T CD4^+^ naïve: (CD3^+^CD4^+^CD45RA^+^CCR7^+^), T CD4^+^ central memory: (CD3^+^CD4^+^CD45RA^-^CCR7^+^), T CD4^+^ effector memory: (CD3^+^CD4^+^CD45RA^-^CCR7^-^), T CD4^+^ EMRA: terminally differentiated effector memory (CD3^+^CD4^+^CD45RA^+^CCR7^-^), T CD8^+^: (CD3^+^CD8^+^), T CD8^+^ naive: (CD3^+^CD8^+^CD45RA^+^CCR7^+^), T CD8^+^ central memory: (CD3^+^CD8^+^CD45RA^-^CCR7^+^), T CD8^+^ effector memory: (CD3^+^CD8^+^CD45RA^-^CCR7^-^), T CD8^+^EMRA: terminally differentiated effector memory (CD3^+^CD4^+^CD45RA^+^CCR7^-^).* LPA: low physically active, HPA: highly physically active, a: significantly different from young, b: significantly different from old HPA, c: significantly different from old LPA. Significant differences were considered p < 0.05. Significance was tested by Independent-Samples Kruskal-Wallis test. Data are presented as mean ± standard deviation (SD).

**S5.** Effects of ageing and physical activity on overall T CD4+ and T CD8+ total puromycin at rest and during activation.

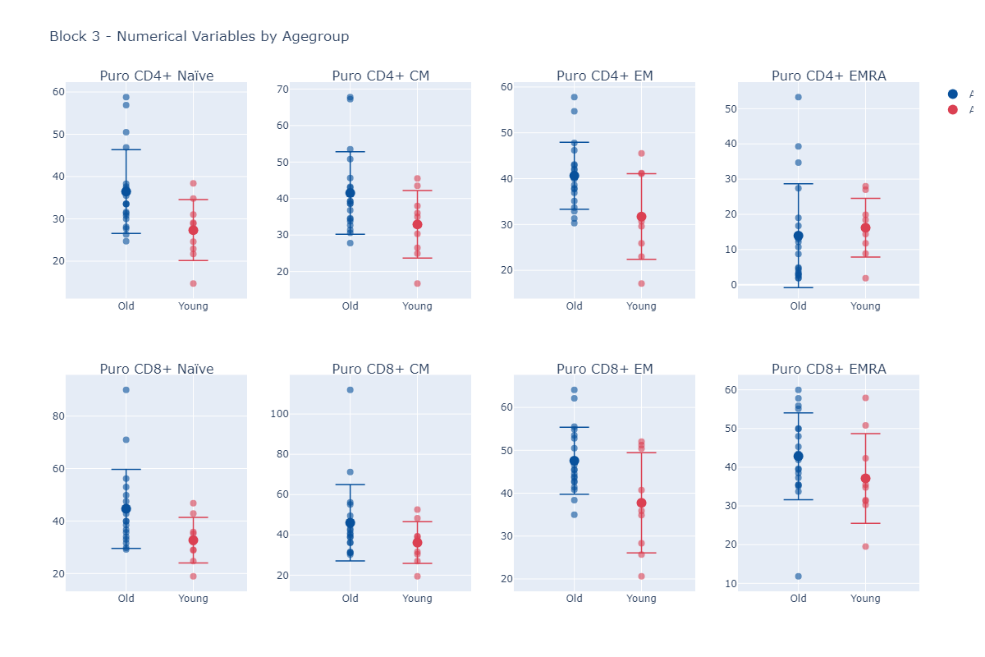

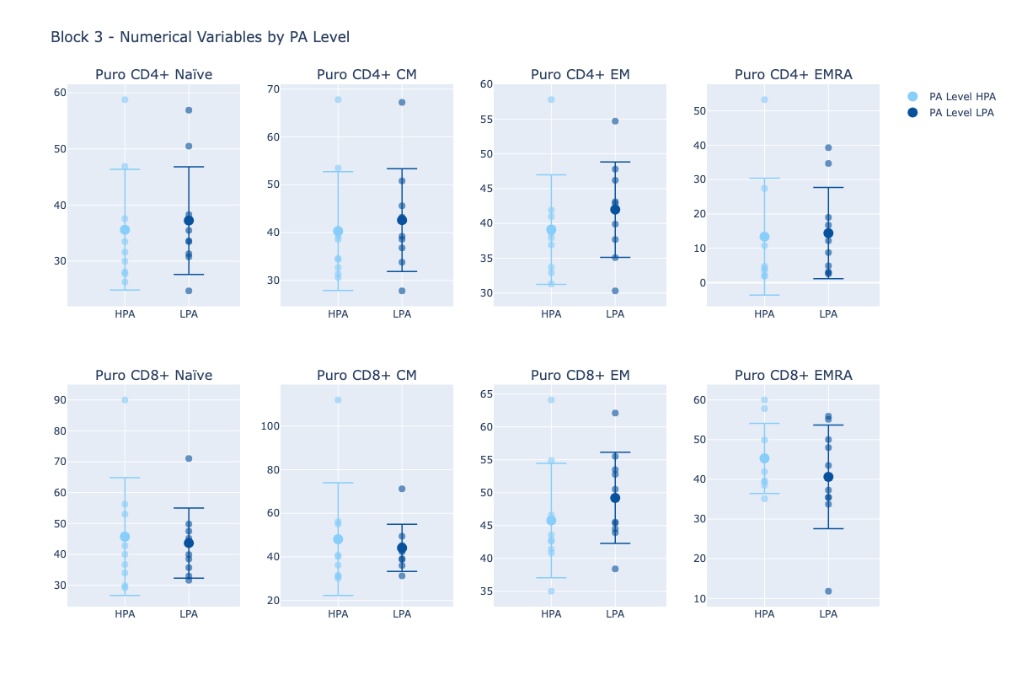

**

**

**

**

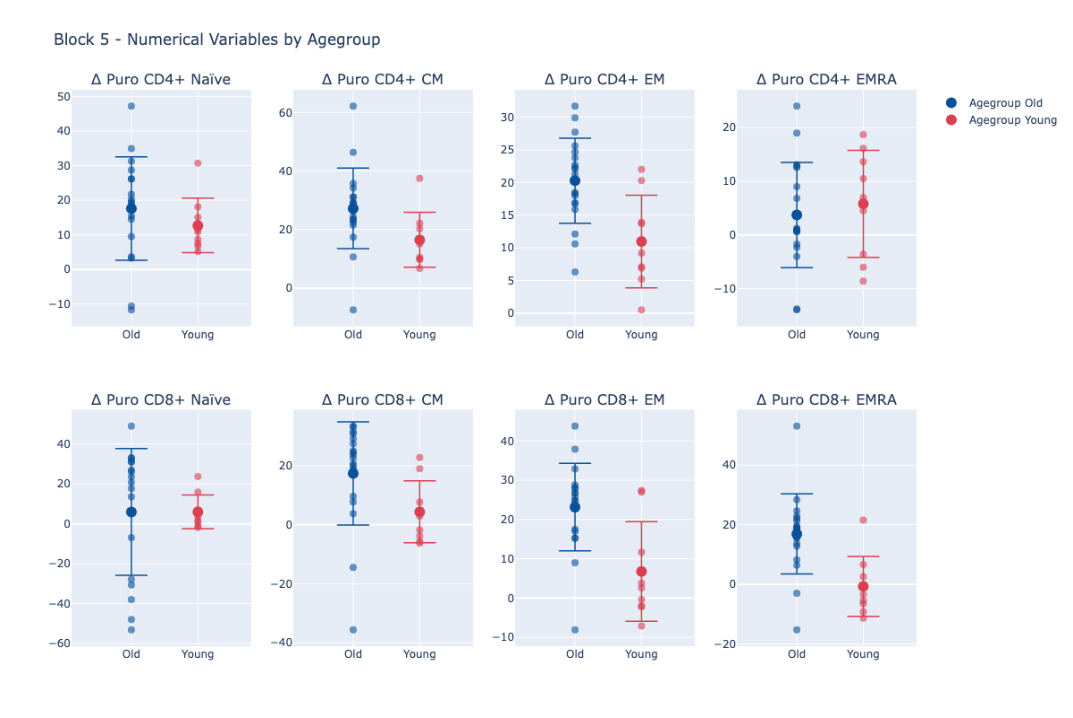

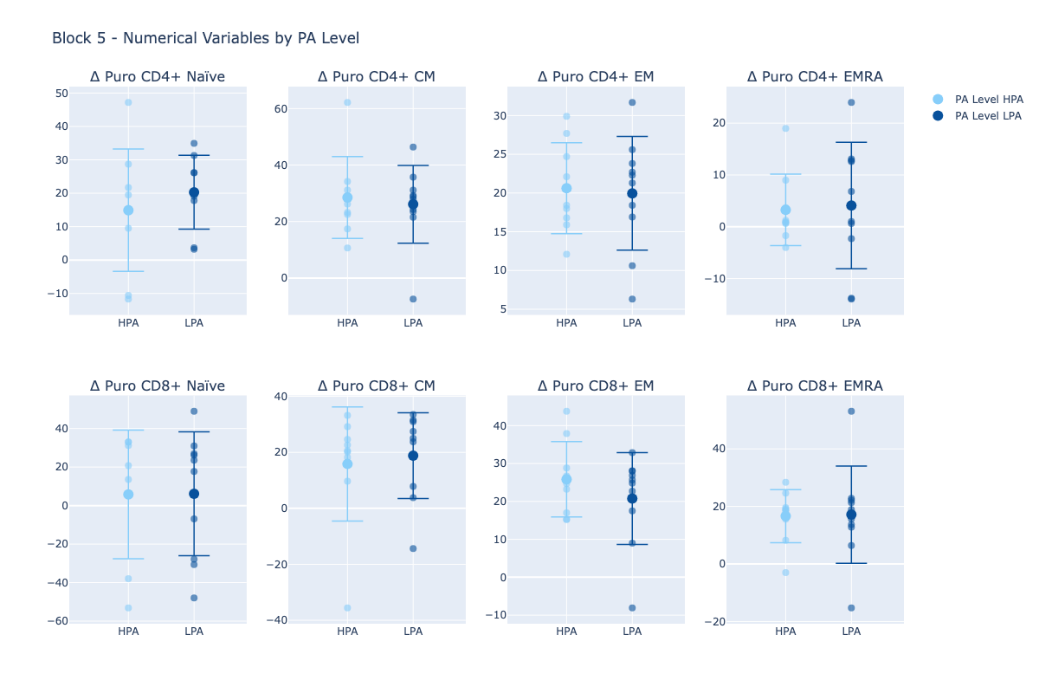

**

**

**

**

**

**Legend:** Puro: total puromycin; Δ: change with stimulation (stimulated minus unstimulated); *: significant difference between age groups (p-value ≤ 0.05).

**S6.** Effects of aging and physical activity on overall T CD4+ and T CD8+ intracellular cytokines at rest and during activation.

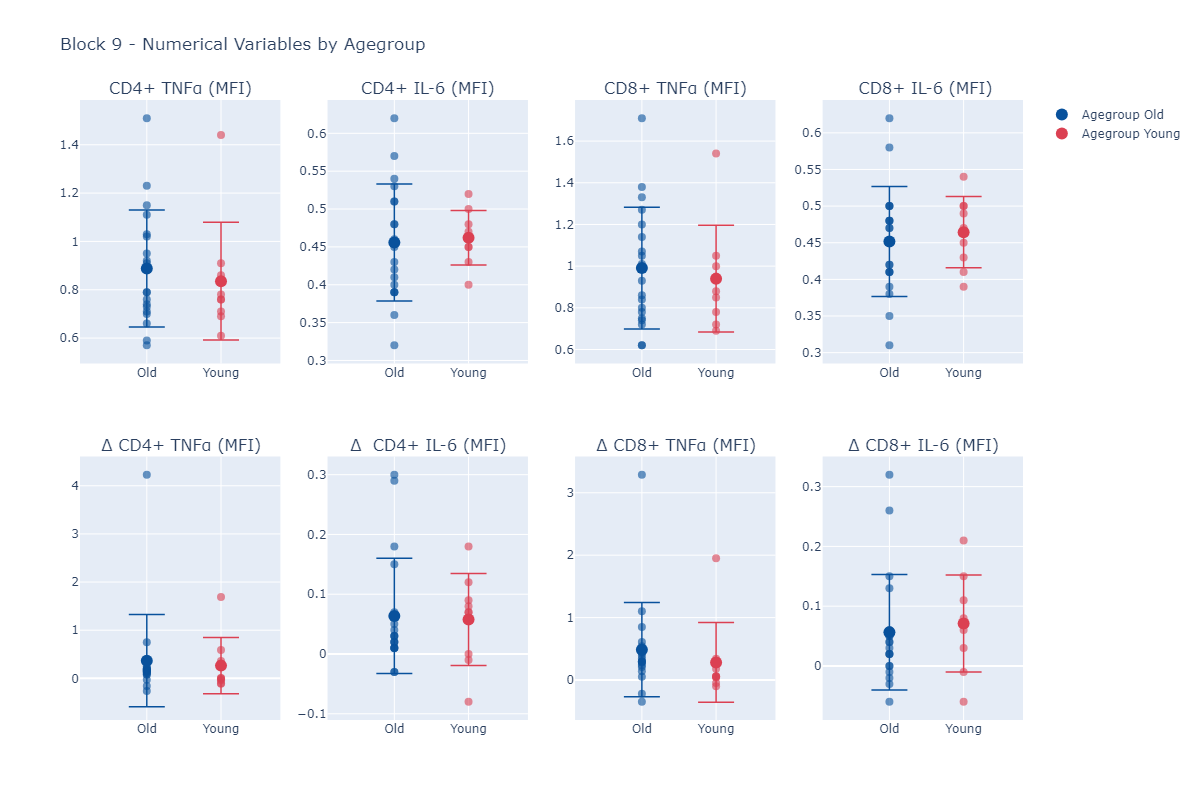

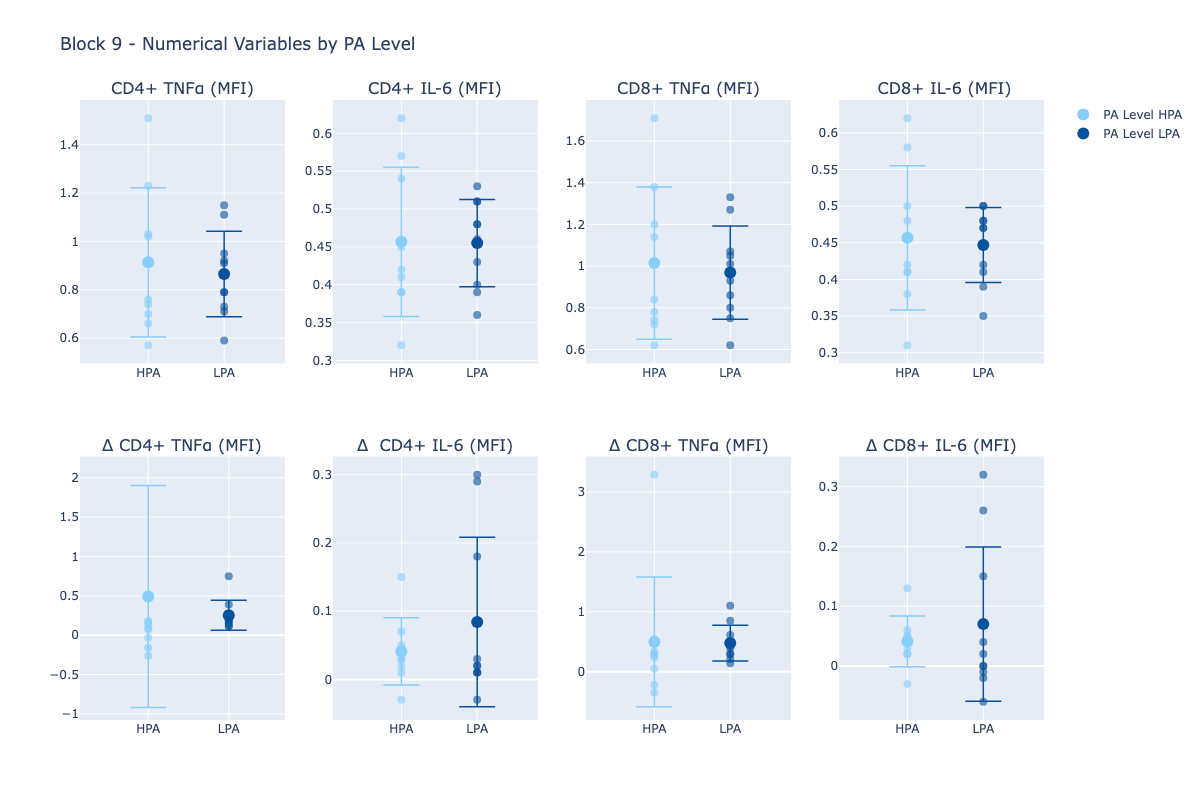

*

**Legend:** GD: Glucose dependence (%); MD: mitochondria dependence (%); Δ: change with stimulation (stimulated minus unstimulated); *: significant difference between age groups (p-value ≤ 0.05).

S7. Effects of aging and physical activity on overall CD4+ and CD8+ metabolic profile.

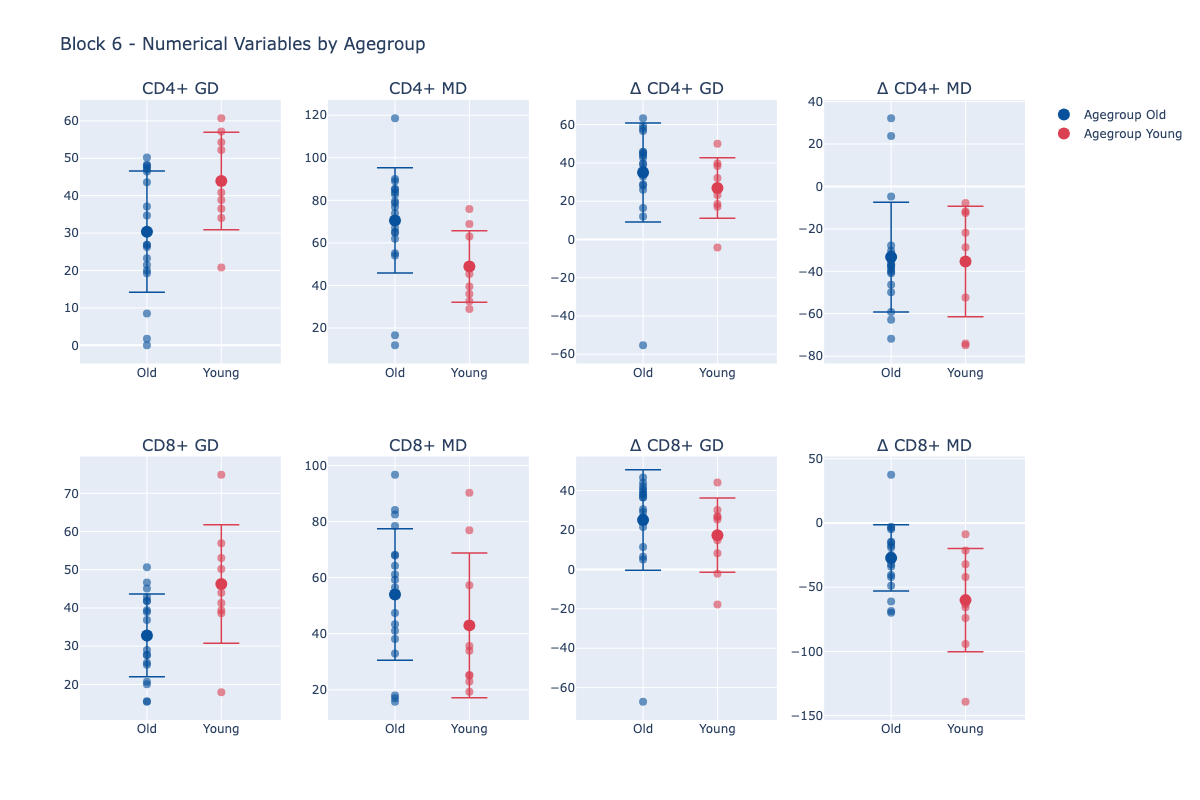

*

*

*

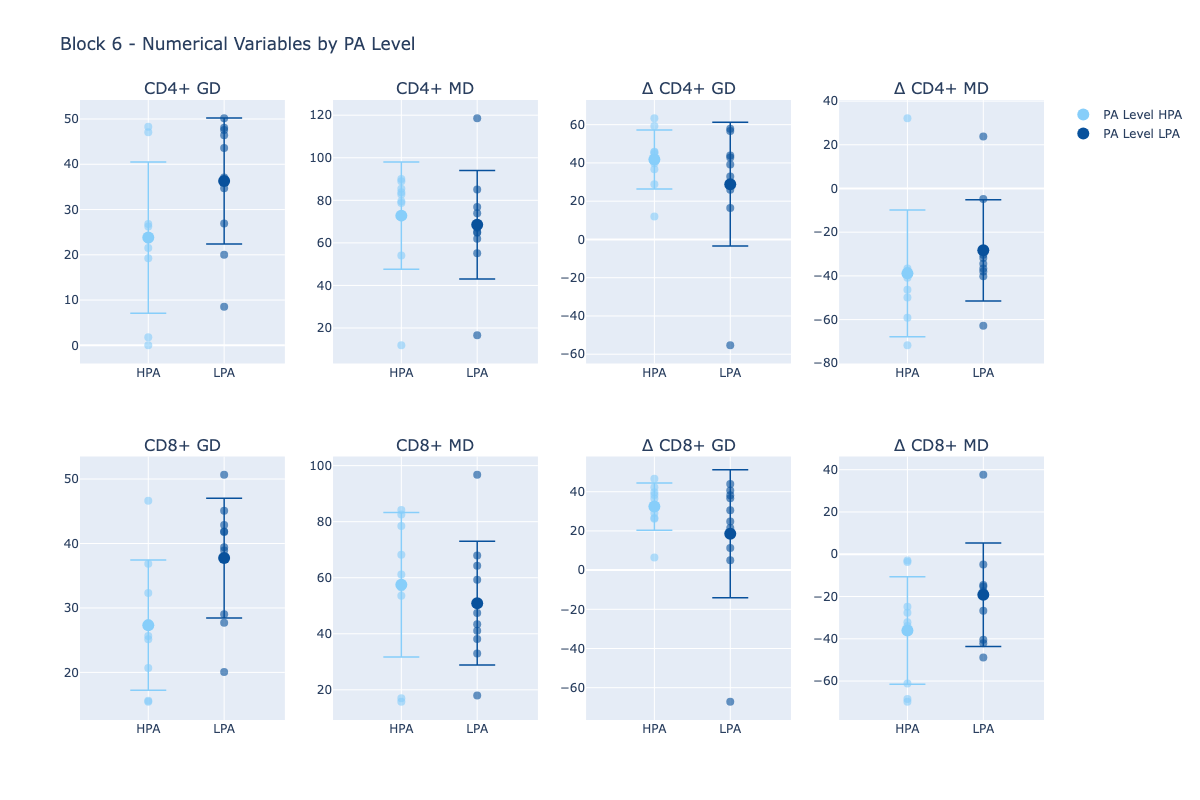

*

**Legend:** GD: Glucose dependence (%); MD: mitochondria dependence (%); Δ: change with stimulation (stimulated minus unstimulated); *: significant difference between age groups (p-value ≤ 0.05).

**S8.** List of key resources.

| **Reagent or Resource** | **Source** | **Identifier** |
| --- | --- | --- |
| ***Antibodies*** | | |
| Alexa Fluor 488 anti-puromycin (Clone: 2A4) | Biolegend | Cat#381506 |
| Alexa Fluor 488 Mouse IgG2a, k isotype Ctrl (Clone: MOPC-173) | Biolegend | Cat#400233 |
| Anti-human CD3 PE Cyanine-7 (Clone: UCHT1) | eBioscience | Cat#25-0038-42 |
| Anti-human CD4 Brilliant Violet 421 (Clone: RPA-T4) | Biolegend | Cat#300532 |
| Anti-human CD8 PE (Clone: QA18A37) | Biolegend | Cat#303804 |
| Anti-human CD8 VioGreen REAfinity (Clone: REA734) | Miltenyi Biotec | Cat#130-110-684 |
| Anti-human CD45RA PerCP (Clone: HI100) | Biolegend | Cat#304156 |
| Anti-human CD197 (CCR7) (Clone: G043H7) | Biolegend | Cat#353214 |
| Anti-human IL-6 PE (Clone: MQ2-13A5) | Biolegend | Cat#501107 |
| Anti-human TNF FITC | BD Pharmingen | Cat#552889 |
| Human TruStain FcX | Biolegend | Cat#422302 |
| ***Chemicals, peptides, and recombinant proteins*** | | |
| autoMACS Running Buffer | Miltenyi Biotec | Cat#130-091-221 |
| Brefeldin A | Merck | Cat#500583 |
| Dimethylsulfoxide (DMSO) | Sigma-Aldrich | Cat#D1435 |
| Ficoll-Paque PLUS | GE Healthcare | Cat#17-1440-03 |
| Fix & Perm Medium A | Invitrogen | Cat#GAS001S100 |
| Fix & Perm Medium B | Invitrogen | Cat#GAS002S100 |
| Heat-inactivated FCS | ThermoFisher | Cat#A3840001 |
| Ionomycin | Merck | Cat#407951 |
| MACSQaunt Running Buffer Concentrate (16x) | Miltenyi Biotec | Cat#130-111-562 |
| Oligomycin from *Streptomyces diastatochromogenes* | Merck | Cat#495455 |
| Peniccilin-Streptomycin | ThermoFisher | Cat#15140122 |
| Phorbel 12-myristate 13-acetate (PMA) | Merck | Cat#524400 |
| Puromycin dihydrochloride from S*treptomyces alboniger* | Merck | Cat#P7255 |
| RPMI Medium 1640 (1X) | Life Technologies | Cat#21875-091 |
| 2-Deoxy-D-Glucose (2DG) | Merck | Cat#25972 |
| 200mM L-Glutamine | ThermoFisher | Cat#25030081 |
| ***Critical commercial assays*** | | |
| FOXP3/Transcription Factor Staining Buffer Set | eBioscience | Cat#00-5523-00 |
| ***Software*** | | |
| FlowJo | FlowJo | v10.10.0 |
| Prism 10 | Graphpad | V8.3 |
| SPSS | IBM | V 29.0 |
